# Pervasive hybridizations in the history of wheat relatives

**DOI:** 10.1101/300848

**Authors:** Sylvain Glémin, Celine Scornavacca, Jacques Dainat, Concetta Burgarella, Véronique Viader, Morgane Ardisson, Gautier Sarah, Sylvain Santoni, Jacques David, Vincent Ranwez

**Author notes:** Equal contribution (Sylvain Glémin, Celine Scornavacca).

## Abstract

Bread wheat and durum wheat derive from an intricate evolutionary history of three genomes, namely A, B and D, present in both extent diploid and polyploid species. Despite its importance for wheat research, no consensus on the phylogeny of the wheat clade has emerged so far, possibly because of hybridizations and gene flows that make phylogeny reconstruction challenging. Recently, it has been proposed that the D genome originated from an ancient hybridization event between the A and B genomes^1^. However, the study only relied on four diploid wheat relatives when 13 species are accessible. Using transcriptome data from all diploid species and a new methodological approach, we provide the first comprehensive phylogenomic analysis of this group. Our analysis reveals that most species belong to the D-genome lineage and descend from the previously detected hybridization event, but with a more complex scenario and with a different parent than previously thought. If we confirmed that one parent was the A genome, we found that the second was not the B genome but the ancestor of *Aegilops mutica* (T genome), an overlooked wild species. We also unravel evidence of other massive gene flow events that could explain long-standing controversies in the classification of wheat relatives. We anticipate that these results will strongly affect future wheat research by providing a robust evolutionary framework and refocusing interest on understudied species. The new method we proposed should also be pivotal for further methodological developments to reconstruct species relationship with multiple hybridizations.

## Introduction

*Aegilops* and *Triticum* form a group of annual Mediterranean and middle east grasses comprising 13 diploid and 18 polyploid species (including durum and bread wheats)^2^ (**Fig. S1**). Most polyploids are allopolyploids^2^. Hybridization is thus possible and has frequently promoted species formation in this group^3^. Moreover, species diversification likely occurred rather rapidly (around 4-7 My^1,3,4^) and some species are highly polymorphic, with a large effective population size^5^. Frequent incomplete lineage sorting (ILS) is thus expected, leading to another source of disagreement between gene and species histories^6^. Both processes could explain why many conflicting results have been obtained for single gene phylogenies so far^7, 8^. In particular, it has proven difficult to resolve the relationships among the three diploid parental donors of the hexaploid wheat, *T. urartu* (A genome), *Ae. speltoïdes* (S genome, considered to be the closest current genome of the B genome), and *Ae. tauschii* (D genome). Recently, based on a multi-gene analysis of the hexaploid wheat and their diploid progenitors, Marcussen et al.^1^ proposed the challenging hypothesis that the D-genome lineage arose 5-6 My ago through a homoploid hybrid speciation between the A-genome and B-genome lineages (A, B and D lineages hereafter), explaining the difficult resolution of a consensual treelike history among these three groups. This result has been questioned and more complex scenarios with several rounds of hybridization have been proposed^2,9,10^. However, none of previous studies included all diploid species and they implicitly referred to the cytology-based classification of wheats, which has not been validated yet by a genome wide analysis.

## Results and discussion

We produced a transcriptome-based dataset of orthologous coding sequences including at least two (and up to four) individuals for each of the 13 diploid *Aegilops/Triticum* species plus one individual of three close outgroups belonging to the Triticeae tribe: *Taeniatherum caput*-*medusae*, *Secale vavilovii* and *Eremopyrum bonaepartis* (**Table S1**). In addition, we used the published sequence of the *Hordeum vulgare* genome (Genome Assembly ASM32608v1) as the most distant outgroup. The transcriptome of each individual was assembled separately and annotated CDSs were stringently clustered and aligned, giving 13288 reliable alignments. After cleaning and processing (see Method), 11033 alignments were retained for the supertree analysis. Among them, the 8739 genes containing at most one sequence per individual were used for the supermatrix analysis and hybrid detection. The 11033 individual gene trees used to construct the supertree with SuperTriplet^11^ were obtained by maximum likelihood (ML) using *RAxML* v8^12^. The total evidence species tree was also obtained by ML from the concatenation of all 8739 one-copy-gene alignments. Both the supertree and the supermatrix approaches gave the same topology (**Fig. 1a**), distinguishing three main clades that we called the A lineage (the two *Triticum* species), the B lineage (*Ae. speltoïdes* + *Ae. mutica*) and the D lineage (all other species), following the simplified terminology of Marcussen et al.^1^. This topology reveals new insights that partly contradict the traditional view of wheat relative evolution. First, while the *Sitopsis* clade (**Fig 1**) is retrieved, *Ae. speltoïdes* is clearly excluded from this clade and appears to be the sister species of *Ae. mutica* (a species whose inclusion in the *Aegilops* genus was partly debated^8^). Second, this topology clarifies what the D lineage of Marcussen et al.^1^ corresponds to by showing that all other diploid *Aegilops* species belong to this clade. Third, it enlightens the relationships among species within the D lineage, while no consensus had emerged so far. Interestingly, the species clustering is in agreement with their geographic proximity, roughly following an east-west distribution (**Fig. S1**).

**Figure 1.**
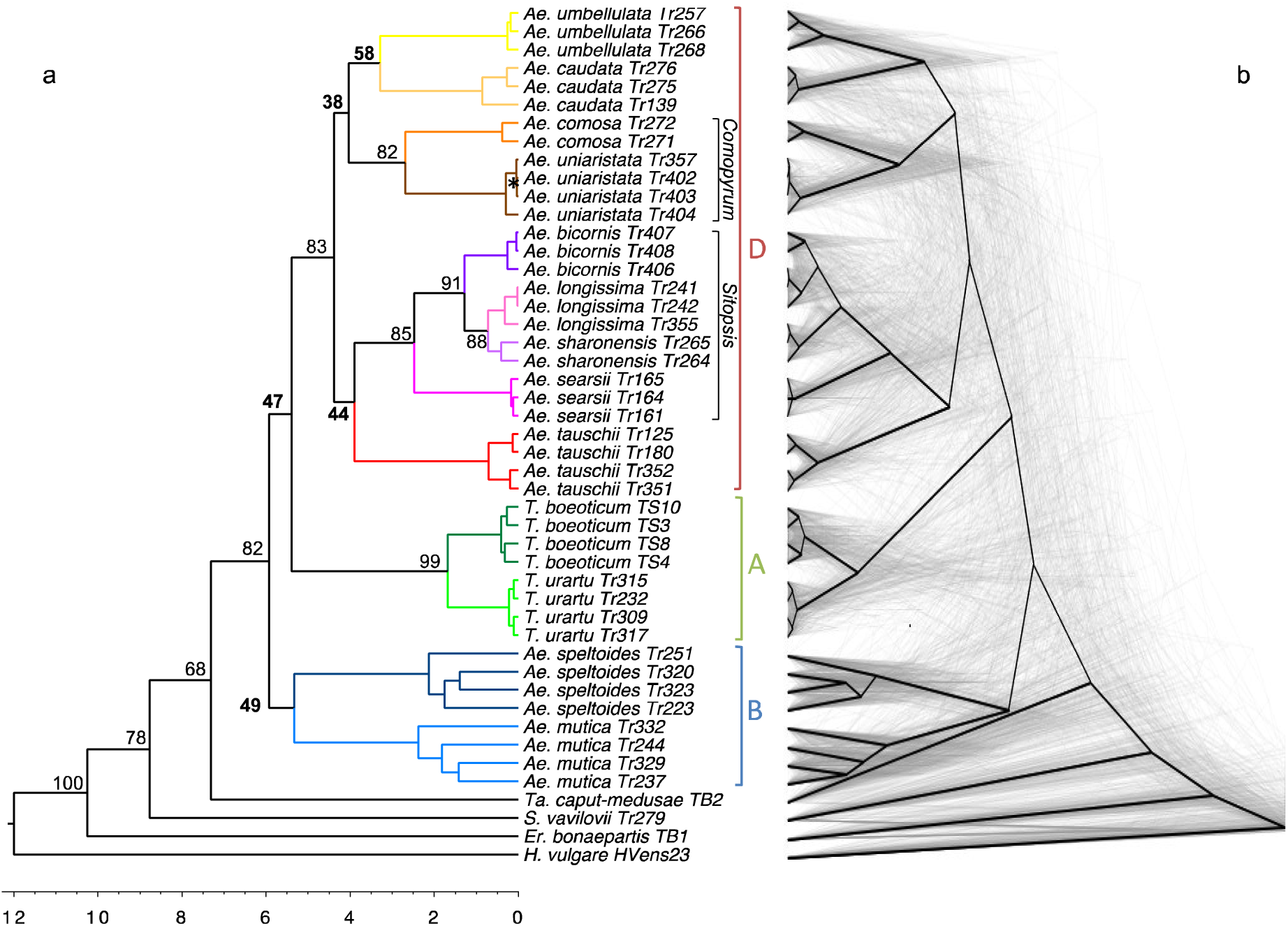
Reconstructed phylogeny of the *Aegilops/Triticum* genus. (**a**) Dated phylogenetic tree of the *Aegilops/Triticum* genus, where each colour corresponds to a species (the same used in **Fig. 1** and **Fig. 3**). This same tree topology was obtained by both the maximum likelihood (ML) analysis of 8739 gene alignment concatenation (supermatrix) and the supertree combination of 11033 individual gene trees. All bootstrap values of the supermatrix analysis are 100 except ^∗^ = 98. Support values for the supertree analysis are given for each inter-species node (% of triplets supporting a given node^11^). Time scale was obtained by making the ML tree ultrametric and assuming a divergence of 12 Myr with *Hordeum*^4^. (**b**) “Cloudogram” of 248 trees (in grey) inferred from non-overlapping 10-Mb genomic windows concatenation. For comparison, the global phylogeny is superposed in black.

However, while the two phylogenomic approaches were fully congruent and the supermatrix tree had very strongly support values (bootstrap = 100 for all but one nodes), the support values of the supertree were low (<60) for 5 over 11 intra-genus nodes (**Fig. 1**) This could be due to both ILS and hybridization. One or more hybridization events have already been proposed but it was difficult to test them directly because previously proposed scenarios invoked ancestral events without considering all extant species. Additionally, current methods developed to jointly infer phylogenetic relationships and hybridization events are not yet able to deal with such large datasets (43 ingroup individuals here), especially with potential nested rounds of hybridization (**Text S1**). As an alternative strategy to detect hybridization in the presence of ILS we proceeded in three steps. First, we searched for all possible hybridization events among triplets of species by counting the number of sites supporting the three possible topologies. Under pure ILS, one major topology and two equivalent minor topologies are expected, while two topologies can predominate over the third one under hybridization^13,14^. From the proportions of the three topologies, a hybridization index can be estimated and tested^1,13,14^. Second, using the phylogeny we obtained as a reference (**Fig. 1**), we identified the possible scenarios compatible with the list of significant hybridization indices (filtered with a very stringent threshold, **Material and methods**). Third, we developed a new likelihood method to discriminate among alternative scenarios involving four taxa and up to two hybridization events (**Texts S2 and S3**). We applied it successively to the groups of species where we identified possible hybridizations.

Hybridization appeared widespread with 40% of triplets showing hybridization index higher than 10%. However, the analysis of triplets composed of two individuals of a same species and a third individual from a second species revealed very low indices, suggesting a nearly absence of recent hybridization (**Fig. 2b**).

**Figure 2.**
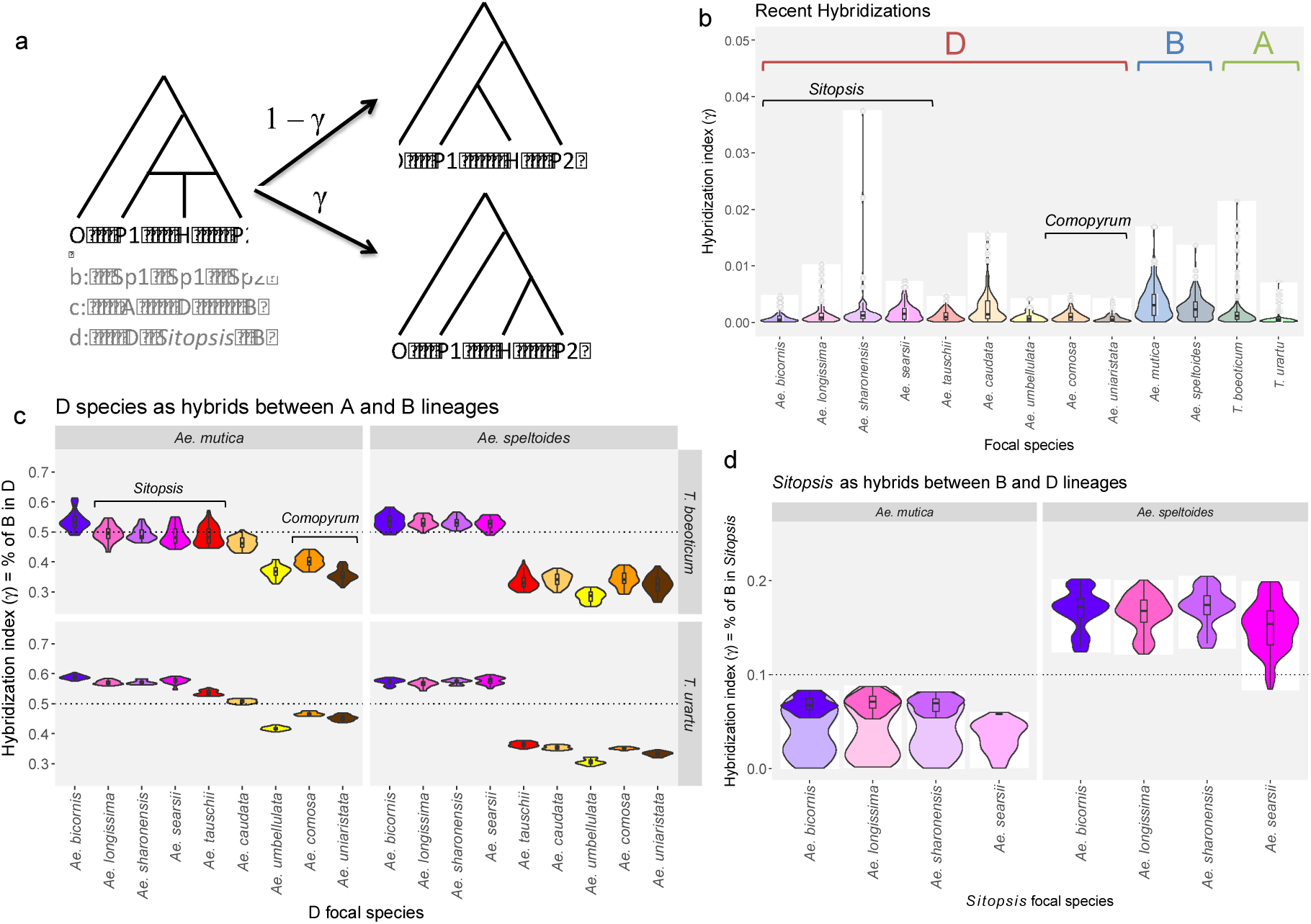
Distribution of the hybridization index among the *Aegilops/Triticum* species. (**a**) For a triplet of species for which a species, H, is a potential hybrid between two parents, P1 and P2, the two possible species trees occur with the proportions 1 – *γ* and *γ*, where *γ* is the hybridization index. The categories of triplets used for the violin plots on figures b, c and d are given in grey. (**b**) Triplets involving two individuals of the same species and a third one from another species. Violin plots are provided for each species being a potential hybrid (focal species), *γ* representing the proportion of gene flow from the other species. Only six indices over 2058 are significantly different from 0 (with the strong threshold *p* < 10^−6^ after Bonferroni correction) and highlighted with bolded dots (**c**) Triplets involving a focal species from the D lineage and parents from the (*T. urartu* or *T. boeoticum*) and B lineages (*Ae. speltoides* or *Ae. mutica*). Violin plots are provided for the nine species of the D lineage as a function of the A and B parents, γ representing the proportion of the B parent. The distributions with *T. boeoticum* as the A parent show higher variance than those with *T. urartu* because of the lower number of available positions to compute γ. The dotted lines correspond to a perfect 50/50 hybridization. All indices are significantly different from 0 (*p* < 10^−6^ after Bonferroni correction) (**d**) Triplets involving *Sitopsis* species as focal species, another D species as one parent and B species as the second parent. Violin plots are provided for each focal *Sitopsis* species *γ* representing the proportion of the B parent. The two A parents (*T. urartu* and *T. boeoticum*) are not distinguished. The dotted line corresponds to 10%. The parts of the distributions corresponding to non-significant indices (*p* < 10^−6^ after Bonferroni correction) are drawn with higher transparency.

In previous studies *Ae. mutica* was not considered as a member of the “B lineage” and the definition of the “D lineage” remained elusive^1,2,15^. Thus, we searched to determine the parental species of the D lineage and whether all species of the D clade descended from the same hybridization event. We considered hybridization indices for triplets where an individual from the D clade could be a hybrid between parents from the A and B clades. The nine species of the D clade show a clear signature of hybridization with a proportion of B species varying from 30 to 70% (**Fig. 2c**), suggesting that all D species are issued from hybridizations between the A and B clades. However, the distribution of hybridization indices was highly heterogeneous, both across potential hybrids and for potential parents, indicating a complex hybridization scenario for the formation of the D clade. Moreover, *Ae. mutica* showed the surprising pattern of being a potential hybrid between *Ae. speltoides* and both A and D species, but at the same time a potential parent of D species (**Text S2**). This can be explained by at least two interwoven hybridization events (**Text S2**). In the most likely scenario (**Fig. 3, Text S3**), *Ae. mutica* – but not *Ae. speltoides* – hybridized with the ancestor of the A clade to give rise to the ancestor of the whole D clade. Before this event, *Ae. mutica* partially hybridized with the ancestor of the A clade. Finally, variation of hybridization indices along chromosomes did not indicate any block structure (**Fig. S2 to S4**) as would be expected under a single event followed by rapid speciation^16,17^. This is hardly compatible with a simple homoploid hybrid speciation scenario and more continuous gene flow may have occurred during species divergence.

**Figure 3.**
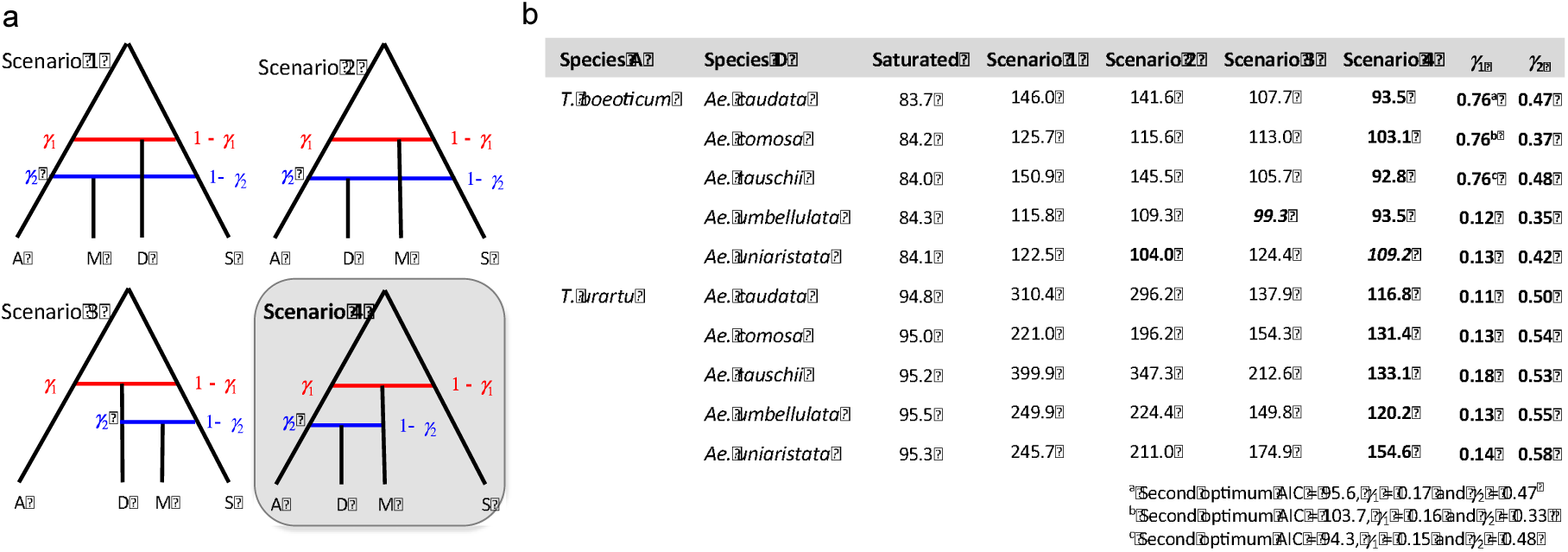
Best scenario for the origin of the D clade determined by the four-taxon maximum likelihood method. (**a**) Schematic representation of the two-hybridization tested scenarios (A: species from the A clade; D: species from the D clade; M: *Ae mutica*; S: *Ae. speltoïdes*). (**b**) AIC of the saturated model and the four tested scenarios. Models were run with the ten different combinations of species from the A and D clades. The best AIC are bolded. In two cases two models have close AIC (the second one is italicized). Scenario 4 is the best model in nine combinations and the second one (with close AIC) in one combination. Point estimates of *γ*_1_ and *γ*_2_ are given for scenario 4: D is the result of two successive hybridization A+S→M then A+M→D. For the three first combinations, there is a second with a very close AIC with a much lower *γ*_1_, in agreement with other values. Scenarios with no or only one reticulations have also been tested and all have much higher AIC (**Supplementary Text 3**).

Among D species, the *Sitopsis* clade showed a distinctive hybridization signature compared to other species (**Fig. 2c,d**). This is compatible with a secondary introgression by *Ae. speltoides* (**Text S2**). This scenario reconciles the morphological and cytological classification of *Ae. speltoides* in the *Sitopsis* clade and some molecular-based phytogenies clearly excluding *Ae. speltoides* from this clade^7,8,18^. This scenario also suggests that chromosome similarities of repetitive elements between *Ae. speltoides* and the *Sitopsis* clade^19,20^ may have resulted of transposable elements exchanges following hybridization, a hypothesis that can now be tested within a clear phylogenetic framework.

Finally, we searched for other possible hybridization events within the D-lineage focusing on triplets where both potential hybrids and parents belong to this clade. We found no signature of hybridization after the divergence of the *Sitopsis* and *Comopyrum* clades, in agreement with the strong supertree supports for these clades (**Fig. 1a**) and with their ancient recognition as taxonomic entities. However, complex patterns were found for *Ae. umbellulata* and *Ae. caudata*, suggesting pervasive gene flow before and during the divergence of this poorly supported clade (**Fig. 1**). *Ae. caudata* showed a particularly strong signature of hybridization, as also suggested by cytogenomics^21^. Although we could not identify the complete scenario, at least two hybridization events are required to explain the results (**Fig. 4, Text S3**).

**Figure 4.**
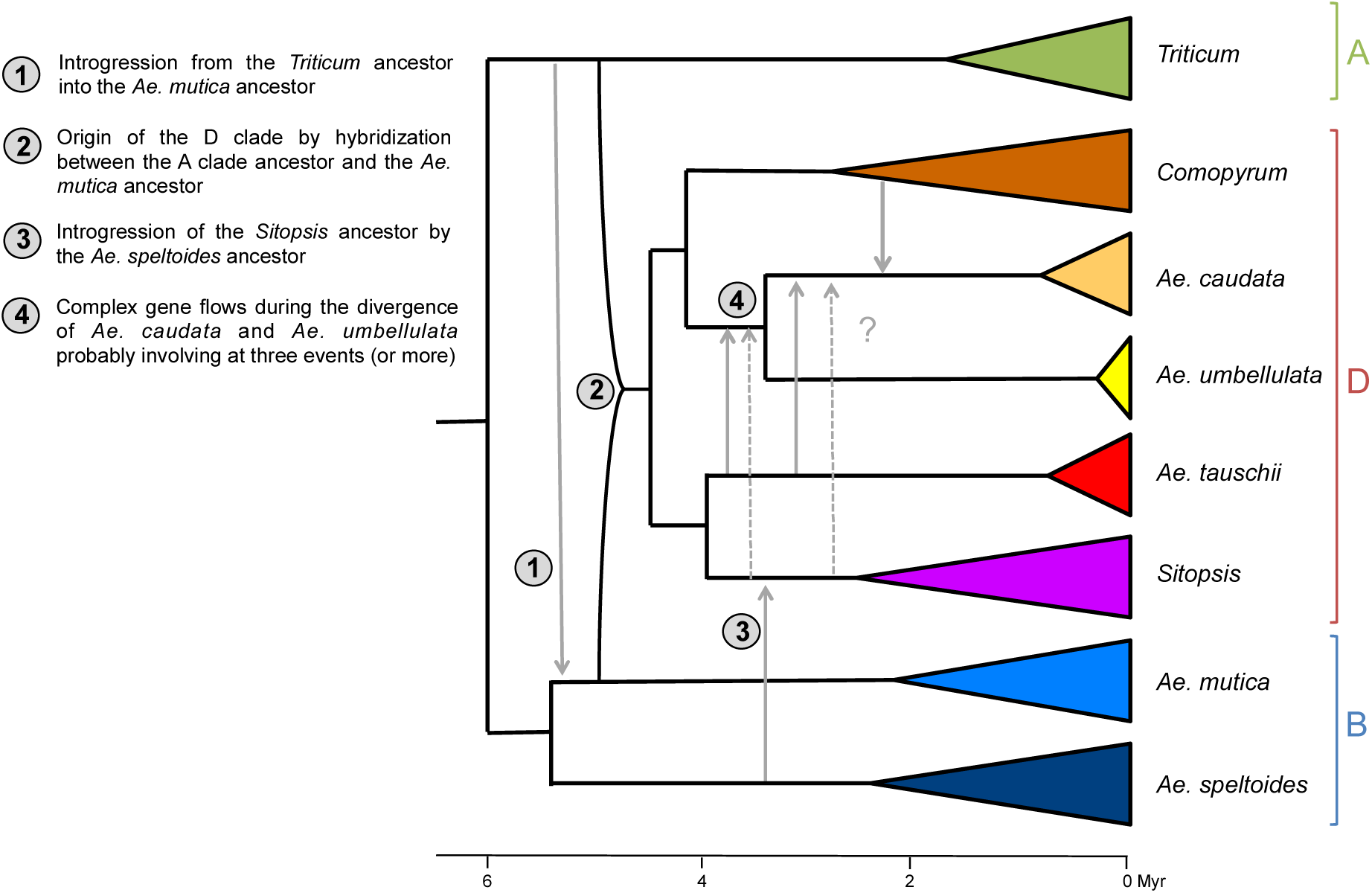
Proposed scenario for the history of diploid *Aegilops/Triticum* species. Proposed hybridization events obtained from the analysis of hybridization indices and from the four-taxa ML method have been added to the global phylogeny with the same time scale (**Fig. 2**). Well-supported clades or individuals from the same species have been collapsed. The length of triangles corresponds to the divergence age as in **Fig. 3**. For event 4, question mark indicates the uncertainties on this complex event. Plain arrows correspond to the most likely detected hybridizations and dotted arrows to possible additional events: we could not exclude hybridization from *Sitopsis* as we could not formally test this hypothesis because of the introgression of *Ae. speltoides* into *Sitopsis* (event 3).

Examples of non-bifurcating speciation histories are accumulating^22-26^. Reconstructing species trees despite ILS and detecting introgression events are now feasible^27^ but inferring the detailed history of multiple and successive events with more than few species remains challenging. Thanks to the development of a new methodological framework, we were able to propose a core reference scenario for the history of diploid *Aegilops/Triticum* species (**Fig. 4**). These results will be pivotal for further research on wheats and their relatives and the approach can be extended to other groups with complex history.

## Material and methods

### Data acquisition

Data were obtained following the same procedure as in Sarah et al.^28^ and redescribed here for comprehensiveness. Sequences of *Triticum boeoticum*, *Taeniatherum caput*-*medusae* and *Eremopyrum bonaepartis* were already obtained by Clement et al.^29^. Sequences of all other species were newly obtained.

All samples were constituted by a combination of leaves (20%) and inflorescence (80%) tissues. RNAs were extracted and prepared separately for each organ and then mixed according to the given proportions. Samples were ground in liquid nitrogen and total cellular RNA was extracted using a Spectrum Plant Total RNA kit (Sigma-Aldrich, USA) with a DNAse treatment. RNA concentration was first measured using a NanoDrop ND-1000 Spectrophotometer then with the Quant-iT™ RiboGreen^®^ (Invitrogen, USA) protocol on a Tecan Genios spectrofluorimeter (Tecan Ltd, Switzerland). RNA quality was assessed by running 1 μL of each RNA sample on RNA 6000 Pico chip on a Bioanalyzer 2100 (Agilent Technologies, USA). Samples with an RNA Integrity Number (RIN) value greater than eight were deemed acceptable according to the Illumina TruSeq mRNA protocol.

The TruSeq Stranded mRNA Library Prep Kit (Illumina, USA) was used according to the manufacturer’s protocol with the following modifications. Poly-A containing mRNA molecules were purified from 2 ug of total RNA using poly-T oligo attached magnetic beads. The purified mRNA was fragmented by addition of the fragmentation buffer and was heated at 94°C in a thermocycler for 4 min. A fragmentation time of 4 min was used to yield library fragments of 250-300 bp. First strand cDNA was synthesized using random primers to eliminate the general bias towards the 3′ end of the transcript. Second strand cDNA synthesis, end repair, A-tailing, and adapter ligation were performed in accordance with the protocols supplied by the manufacturer. Purified cDNA templates were enriched by 15 cycles of PCR for 10 s at 98°C, 30 s at 65°C, and 30 s at 72°C using PE1.0 and PE2.0 primers and the Phusion^®^ High-Fidelity PCR Master Mix (Thermo Fischer Scientific, USA). Each indexed cDNA library was verified and quantified using a DNA 100 Chip on a Bioanalyzer 2100 to build pooled libraries made of twelve, equally represented, genotypes.

The final pooled library was quantify by qPCR with the KAPA Library Quantification Kit (KAPA Biosystems, USA) and provided to the Get-PlaGe core facility (GenoToul platform, INRA Toulouse, France http://www.genotoul.fr) for sequencing. Each final pooled library (12 genotypes) was sequenced using the Illumina paired-end protocol on a single lane of a HiSeq3000 sequencer, for 2 × 150 cycles.

### Transcriptome assembly and annotation

Reads have been cleaned and assembled following the pipeline described in Sarah et al.^28^ and recalled here for comprehensiveness. Reads were preprocessed with cutadapt^30^ using the TruSeq index sequence corresponding to the sample, searching within the whole sequence. The end of the reads with low quality scores (parameter -q 20) were trimmed and we only kept trimmed reads with a minimum length of 35 bp and a mean quality higher than 30. Orphan reads were then discarded using a homemade script. Remaining paired reads were assembled using ABySS^31^ followed by one step of Cap3^32^. Reads returned as singletons by the first assembly run were discarded. Abyss was launched using the paired-end option with a kmer value of 60. Cap3 was launched with the default parameters, including 40 bases of overlap, and the percentage of identity was set at 90%.

We slightly modified the *Rapsearch* program^33^ to make its blast formatted output compatible with the expected input format of prot4est. We used this modified version of Rapsearch to identify protein sequences similar to our contigs in either plant species of Uniprot swissprot (http://www.uniprot.org) or in the Monocotyledon species of greenphyl (http://www.greenphyl.org/cgi-bin/index.cgi). We then used the *prot4est* program^34^ to predict the CDS embedded in our contigs based on the following input: *Rapsearch* similarity output, *Oryza* matrix model for de-novo based predictions and the codon usage bias observed in *T. boeoticum.*

Short sequences are often difficult to cluster into reliable orthologous groups and are not very informative for phylogeny inference; we hence discarded predicted CDS with less than 250bp as done in a similar context to populate the OrthoMaM database^35^. The total numbers of contigs per species is given in **Table S2.**

### Orthologous search

We relied on *usearch* v7^36^ to cluster the predicted CDSs. We designed a four-step approach that limits the impact of taxon sampling and sequence ordering during cluster creation, avoids assigning sequences to an arbitrary cluster in case of tile and can easily handle our large dataset (both in terms of required memory and computation time). First, for each species of the ingroup, we selected the accession with the highest number of CDSs to represent this species during the first step of cluster creation. Second, we used *uclust* to cluster these sequences and to output the median sequence of each cluster, which will be used as cluster bait. Third, we used *usearch* to identify, for each predicted CDS, the set of clusters for which the considered CDS and the cluster bait had a similarity above 85% along at least 50% of their length. Finally, all predicted CDSs having such a similarity with one single cluster bait were assigned to this cluster; all others were discarded.

### Alignment and cleaning

Following the strategy used to populate the OrthoMaM database^35^, CDSs were aligned at the nucleotide (NT) level based on their amino acid (AA) translation combining the speed of *MUSCLE*^37^ and the ability of *MACSE*^38^ to handle sequence errors in predicted CDSs resulting in apparent frameshift and erroneous amino acid translations. In more detail, for each cluster we did the following: First, CDSs were translated into AA, these AA sequences were then aligned using *MUSCLE*, and the obtained protein alignment was used for deriving the nucleotide one using *MACSE reportGapsAA2NT* routine. Second, this nucleotide alignment was refined using *MACSE refineAlignment* routine. Finally the resulting amino acid alignment was cleaned with *HMMcleaner*^39^ and a homemade script (that will be part of the next MACSE release) was used to report the obtained amino acid masking at the nucleotide level.

### Phylogeny reconstructions

Gene trees were inferred with *RAxML* v8^12^ using the GTR model with a 4 categories gamma distribution (GTR+T4) to accommodate for evolution rate heterogeneity among sites, and using *RAxML* fast-bootstrap option (-f).

BppReroot of the BppSuite^40,41^ was used to reroot the 13288 gene trees, using as outgroups the following ordered list of species: *H*_*vulgare*, *Er*_*bonaepartis*, *S*_*vavilovii* and *Ta*_*caputMedusae.* In more detail, for each of the gene tree, we considered each species of the outgroup list one after the other until finding the first one present in the current gene tree (if none was found we discarded the tree). Having identified the most relevant outgroup species for this gene tree, we then checked whether all the individuals of this outgroup species formed a monophyletic clade; if yes, we rooted the tree on this clade, otherwise we discarded the tree. This resulted in a forest of 12959 rooted gene trees, which we denoted by ***F_i_***.

Since our aim here was to build a phylogeny of species and not of individuals, we focused on the identification of reliable species clades from the information contained in the gene trees. Therefore, we derived from ***F_i_*** two forests of multi-label trees by renaming each sequence by the species to which it belongs to (forest ***F_m_***) and keeping only clades with a bootstrap value greater than 95 (forest ***F^95^_m_***). Almost all trees (99,99%) in these forests are multi-labelled as alignments include several individuals for at least some species. We thus used *SSIMUL*^42^ to process the multi-label of trees of ***F_m_*** and ***F^95^_m_*** by turning —without losing phylogenetic signal when possible —its multilabelled trees into single-labelled trees. This was done by removing a copy of each pair of isomorphic sibling subtrees^42^. We denoted by ***F_s_*** and ***F^95^_s_*** the new forests obtained by pruning isomorphic trees of ***F_m_*** and ***F^95^_m_***, respectively. We used *SuperTriplets*^11^ to construct a supertree from the 11033 trees in ***F^95^_S_***. The resulting supertree is depicted in Fig.2. The support values given by SuperTriplets to the clades are very low (only three clades have a support greater than 90); this shows that, even if we only keep clades with a support greater than 95, ***F^95^_s_*** contains a high level of contradiction.

### Supermatrix analysis

In the forest ***F_i_***, some trees are also multi-labelled at the individual level either because of paralogy or because the two allelic copies were split. From ***F_i_***, we extracted the set of 8739 trees containing at most one sequence per individual. We built the concatenation of all the 8739 alignments corresponding to these trees, giving a supermatrix with one sequence for all individuals. We inferred the phylogeny from this supermatrix with RAxML v8^12^ using the GTR+Γ4 model and the fast-bootstrap option. The resulting phylogeny has the same topology than the supertree shown in Fig.2 and all nodes but one have bootstrap values equal to 100. Using the *Hordeum* genome as a reference, we also concatenated genes in 10-Mb windows along chromosomes, obtaining 298 alignments with at least three genes per window. (For this analysis, 5976 genes were kept since the others could either not be assigned to a position on the *Hordeum* genome or were isolated—one or two sequences—in their 10-Mb window). We reconstructed the phylogeny of each alignment using the same method. The global tree and the 298 10-Mb trees were made ultrametric using the *chronos* function of the *ape* R package^43^. Among the 298 10-Mb trees, only 248 contained all individuals. We used them to draw the “cloudogram” presented in Figure 2b using *Densitree*^44^.

### Detection of hybridization events

We used the same supermatrix alignments to detect possible hybridization events by applying the rationale developed by Meng and Kubatko^13^ and Kubatko and Chiffman^14^. Note that this was also the rationale used to propose the hybrid origin of the D genome^1^. In broad strokes, if we consider a triplet of lineages, A, B and C, with B being a hybrid between A, in proportion 1 – *γ*, and C, in proportion *γ*, then the probabilities of the three rooted topologies are given by:

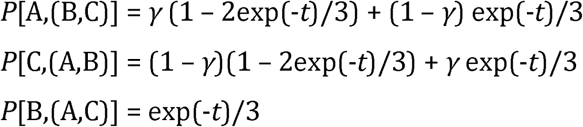

where *t* represents the time between speciation events on the parental trees measured in 2*N_e_* generations. It can be easily shown that:

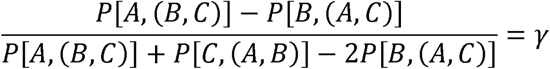

Note also that 2*P*[B,(A,C)] = 2exp(-*t*)/3 directly gives the probability of incongruence due to ILS^6^. *γ* can thus be estimated by counting the number of two-state (*i* and *j*) positions supporting each topology, using an outgroup (O) to polarize mutations^14^. Considering the order O/A/B/C, we have:

‐ *x = #i*,*i*,*j*,*j* → A,(B,C)
‐ *y = #i*,*j*,*j*,*i* → C,(A,B)
‐ *z = #i*,*j*,*i*,*j* → B,(A,C)

So we can define a hybridization index that is an estimator of *γ*:

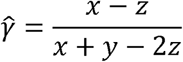

To test the significance of this estimator, that is to identify *γ* values not due to random sampling under pure ILS, Kubatko and Chiffman^14^ proposed a statistics (called the “Hils statistics”) that is normally distributed with mean zero and variance one. It allows rapidly detecting significant potential hybrids among all possible triplets in a large phylogeny. We used this test to filter out the *γ* estimates and only consider significant ones. Because of the high rate of false positive of this test^14^ and of the large number of sites in the alignment we used the very stringent threshold of 10^−6^ (instead of 0.05) after Boferroni correction for multiple testing. In addition we focused on major events for which *γ* > 10%. It is also worth noting that the above rationale implicitly assumes that the effective size, *N_e_*, remained the same in the two diverging A and C lineages. Relaxing this assumption bias the estimation of *γ*, but *ŷ* is still expected to be null only without hybridization, so that detection of hybridization is conservative. However, a single *ŷ* value can be difficult to interpret when multiple hybridization events occurred. Thus, we first computed the statistics for all triplets to list all possible hybridization events. Then we tested formally the proposed scenarios within a ML framework (see below).

To compute the values of *ŷ* for each triplet of individuals, we applied a modified version of the Hyde program^14^ to allow retrieving the counts of each patterns *x*, *y*, *z* and not only the Hils statistics. As outgroup we used the consensus sequence of the four outgroup species in order to limit homoplasy, which can bias statistics. For each triplet, we ordered topologies and species such that *x* > *y* > *z* and computed *ŷ*. We applied it to the full alignment and to the 298 10-MB window alignments.

With 43 ingroup individuals, 74046 triplets are possible, making the analysis of individual triplets useless. Instead, we parsed results hierarchically based on the clades previously obtained with phylogenetic analyses: we started from triplets of species belonging to the same species and sister species until triplets of species belongings to the three main clades (A,B,D), From this analysis (detailed in **Text S2**), we proposed a series of hybridization scenarios. To detect possible heterogeneity of ILS and hybridization events across the genome, we also analysed variation of the two statistics along chromosomes and performed simulations to evaluate the size of hybridization blocks across the genome (see **Text S4**).

### Test of multiple hybridization scenarios

With three taxa, only three rooted topologies are possible, leaving only two degrees of freedom to estimate scenario parameters, which is not sufficient if multiple hybridization events occurred. Using four taxa, ten informative bi-allelic sites patterns are possible leaving nine degrees of freedom to infer scenarios (**Text S3**). Noting 0 the ancestral and 1 the derived allele, the ten informative site patterns are: 0|0111, 0|1011, 0|1101, 0|1110, 0|0011, 0|0101, 0|0110, 0|1001, 0|1010, and 0|1100. Scenarios with four taxa and up to two hybridization events can be described with eight parameters (see below and **Fig. S2.3**). In **Text S3**, we show how to write the probabilities of the ten site patterns under a four-taxon multi-species coalescent model with up to two hybridization events. To do so we need to compute both the probabilities of the compatible gene tree topologies and the expected length of branches for which the occurrence of a mutation leads to the given pattern. Then, to obtain the probabilities for a full scenario we need to take the weighted sum of all possible gene trees embedded in the four-taxon hybridization network. Formally the probability of site pattern *i* within a scenario 𝕊 can be written as:

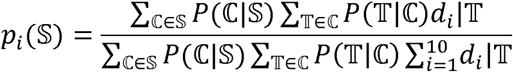

where ℂ is a component of the decomposition of the scenario 𝕊 (species tree or one-reticulation network, see below), 𝕋 is a gene tree embedded in component ℂ, and *d_i_*|𝕋 is the expected length of the branch where a mutation leads to site pattern *i* for a given gene tree, 𝕋.

Scenarios with two **non-nested** reticulations can be decomposed into the four *trees displayed* by the corresponding phylogenetic network^45,46^. We first obtained the vectors of expected branch lengths leading to the ten site patterns for these four trees – denoted by ***I_i_***, with i ranging from 1 to 4. Note that the longer a branch the higher the probability for a mutation to occur so that branch lengths directly impact observed pattern frequencies. We hence enumerated all possible gene trees embedded in a given four-taxon species tree and computed both the probabilities of the compatible topologies and the mean length of the branches where the occurrence of a mutation leads to a given site pattern. Probabilities and branch lengths are function of divergent times and coalescent rates (**Supplementary Text 3**). Then a full scenario with hybridization can be obtained by combining the corresponding trees with their respective weights. Consider two non-nested hybridization events with proportions of the parental lineages being *γ*_1_ and 1 – *γ*_1_ for the first event and *γ*_2_ and 1 – *γ*_2_ for the second one. The vector of probabilities for the full network is thus:

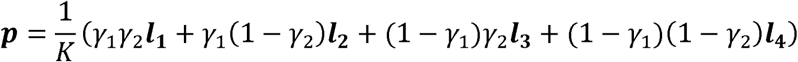

where *K* is a normalization constant such that 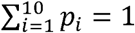

For scenarios with two **nested** reticulations, hybridization and coalescent processes cannot be fully decoupled^46^, and some embedded coalescent trees must be computed directly on a network component instead of a tree component. If only one species is issued from two nested hybridization events (the only case considered here), the initial network can be decomposed into two trees in proportions *γ*_1_*γ*_2_, (1 – *γ*_1_)*γ*_2_ and one one-reticulation network in proportion (1 – *γ*_2_). Noting ***I*_1_** and ***I*_2_** the vectors of branch lengths for the two trees and ***λ*** for the one-reticulation network, the vector of probabilities for the full network is thus:

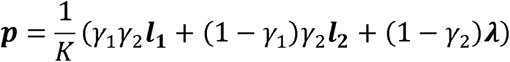

where *K* is the normalization constant.

Noting **v** the vector of the number of positions corresponding to the ten bi-allelic patterns, the likelihood of a network is given by the multinomial sampling:

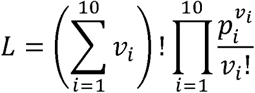

By fixing either *γ*_1_ or *γ*_2_ to 0 or 1 we obtain a scenario with only one reticulation and fixing both parameters to 0 or 1 a tree-like scenario without any reticulation. A scenario with one reticulation has six free parameters and without any reticulation only four. As all scenarios cannot be nested in each others we used Akaike Information Criterium (AIC) to compare them where AIC = 2*k* – 21n(*L*). Below we show how to compute the ***p*** vectors. Likelihood maximization was made with a *Mathematica* script provided in **Supplementary File 1.** The FindMaximum function was used with ten random starting points to ensure convergence to the global optimum.

In the following, we excluded the *Sitopsis* clade from the analyses because of the additional hybridization with *Ae. speltoides.* We first applied the model to the four taxa: A clade, **D** clade, *Ae. mutica* and *Ae. speltoides* to elucidate the origin of the **D** clade. Because the triplet analysis showed heterogeneity among species we run the model successively for the ten combinations of the two species from the A clade (*T. boeoticum* and *T. urartu*) and the five species of the D clade (*Ae. caudata*, *Ae. comosa*, *Ae. tauschii*, *Ae. umbellulata* and *Ae. uniaristata*). As only four sequences are required for this analysis we used the strict consensus of the different sequences of a same species. As for the triplet analysis we used the consensus sequence of the four outgroup to polarised mutations. We only tested scenarios where the D clade and *Ae. mutica* could be potential hybrids as there was no signature that neither *Ae. speltoides* nor the two *Triticum* species could be potential hybrids according to the distribution of hybridization indices. We then applied the method to *Ae. caudata*, *Ae. tauschii*, *Ae. umbellulata* and either *Ae. comosa* or *Ae. uniaristata* from the *Comopyrum* clade.

### Pattern of hybridization indices along chromosomes

Under the hypothesis of a simple and single hybridization event, blocks of parental genomes should be detectable along chromosomes^16,17^. Here, the corresponding signature should be an alternation of hybridization index close to 0 and close to 1. We focused on the hybrid origin of the **D** lineages and re-computed the hybridization index with (A, *Ae. mutica*, D) triplets for every 298 10-Mb concatenations alignments obtained for drawing the cloudogram on **Fig. 2.** In D, we excluded the *Sitopsis* clade that has likely experienced a secondary introgression event from *Ae. speltoides.* We then took the mean of the index for each 10-Mb window. The index distribution is roughly bell-curved with no extreme values (0 or 1) (**Fig. S2**) without any specific pattern along chromosomes (**Fig. S3**). This shows that there is no large block of pure A or B origin in D genomes, as it would be expected under a simple homoploid hybrid speciation scenario^16,17^.

To confirm this conclusion we ran simple simulations. We drew a series of consecutive genomic segments in a Poisson distribution with different mean and alternatively attributed values 0 (A origin) or 1 (B origin) to these segments. Then the hybridization index was computed as the mean over 10-Mb windows. **Fig. S4** clearly shows that the mean size of such blocks should be lower than the chosen 10-Mb windows size.

## Acknowledgements

This publication is the contribution ISEM XX of the Institut des Sciences de ľEvolution de Montpellier (UMR 5554 – Université de Montpellier-CNRS-IRD-EPHE). This work was supported by the French Centre National de la Recherche Scientifique and Agence Nationale de la Recherche (ANR-11-BSV7-013-03). Analyses were performed on the computing cluster of the Montpellier Bioinformatics Biodiversity (MBB) platform, the South Green Bioinformatics platform and on the UPPMAX platform in Uppsala.

## Author contributions

‐ Conception: JDav, SG, VR
‐ Funding acquisition: JDav, SG
‐ Biological data acquisition and management: CB, W
‐ Sequence data acquisition: GS, MA, SS
‐ Method development: SG
‐ Data analysis: CS, JDai, SG, VR
‐ Interpretation of the results: CB, CS, JDav, SG, VR
‐ Drafting the manuscript: SG
‐ Reviewing and editing the manuscript: CB, CS, JDav, SG, SS, VR

## Materials and correspondence

Sylvain Glémin

## Supplementary Figures

**Figure S1.**
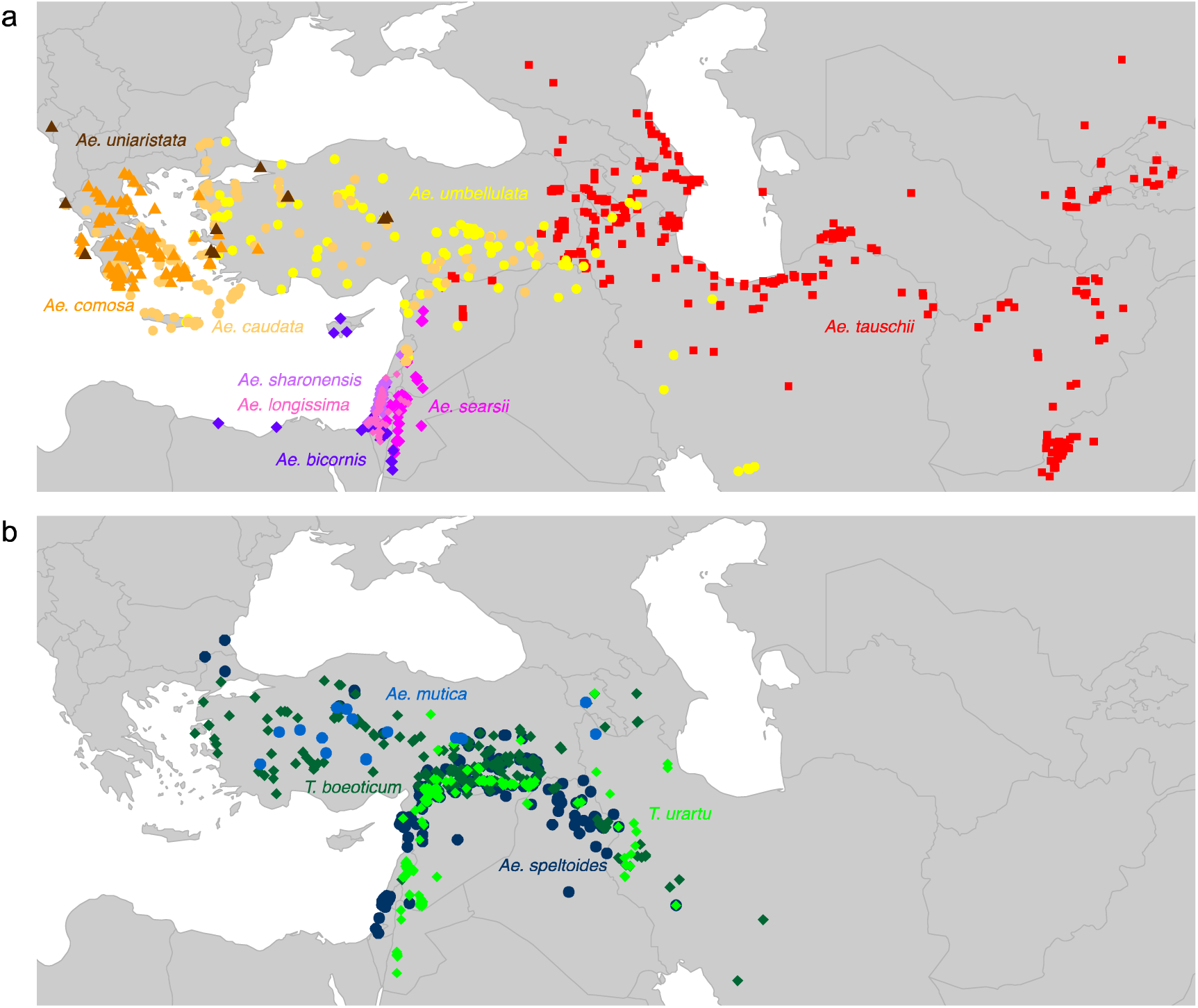
Geographic distribution of the 13 diploid *Aegilops/Triticum* species. (**a**) Nine species belonging to the D lineage (see **Fig. 2**) (**b**) Species belonging to the A and B lineages. Each dot corresponds to an observation retrieved from GBIF (http://www.gbif.org).

**Figure S2.**
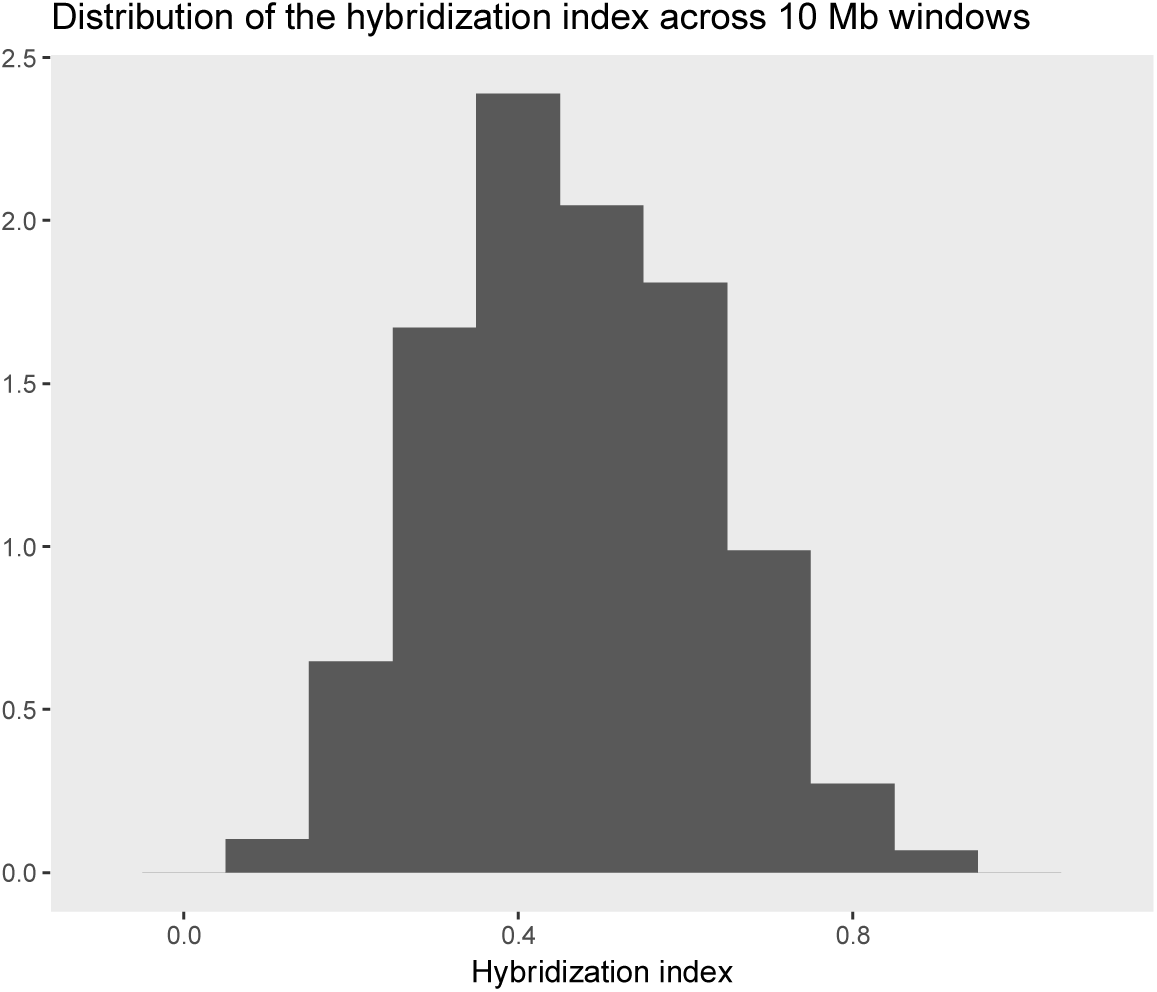
Distribution of the mean hybridization index for (A, *Ae. mutica*, D) triplets (proportion of *Ae. mutica* in D) across 10-Mb concatenations.

**Figure S3.**
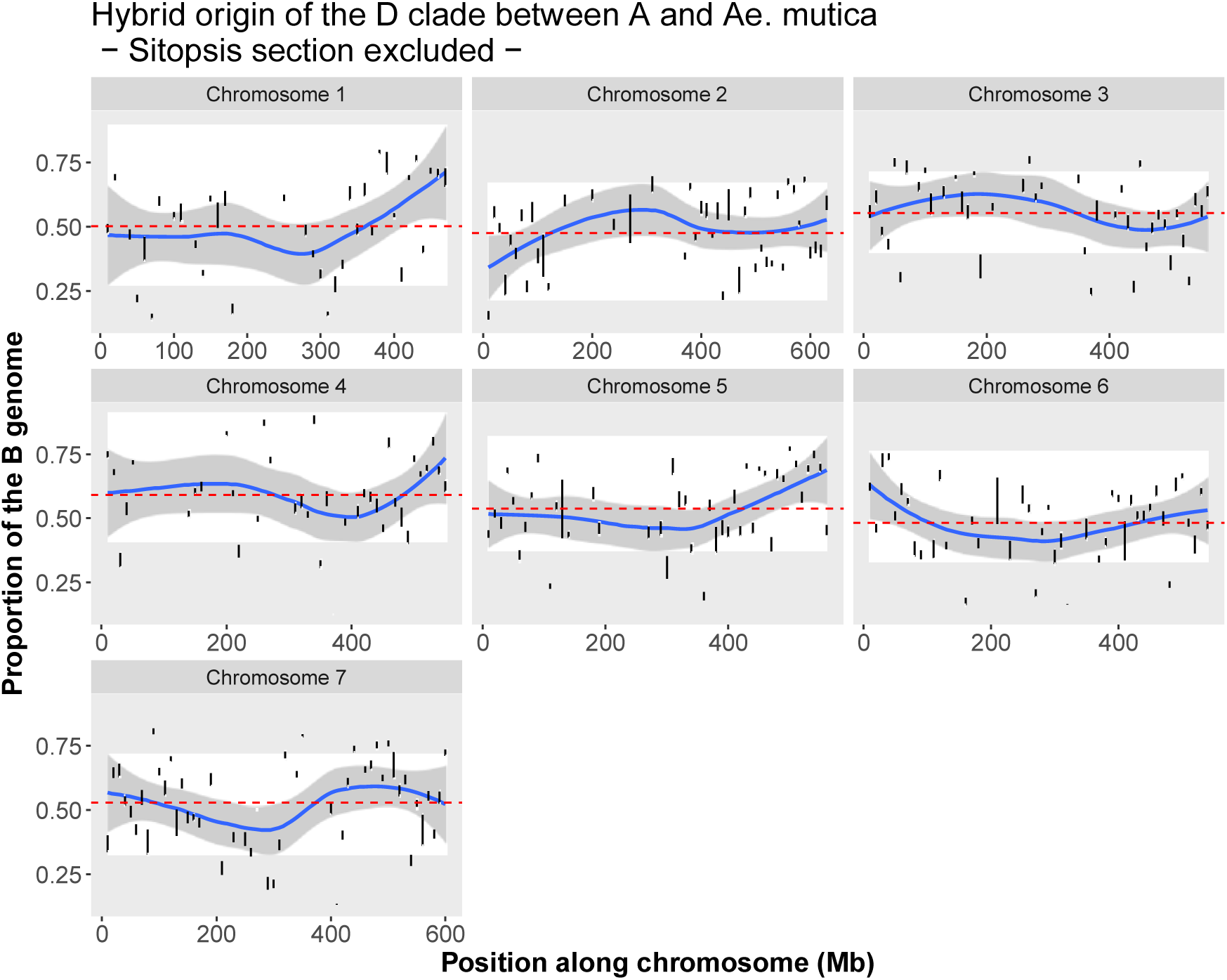
Distribution of the mean hybridization index for A, *Ae. mutica*, D triplets (proportion *of Ae. mutica* in D) along chromosomes.

**Figure S4.**
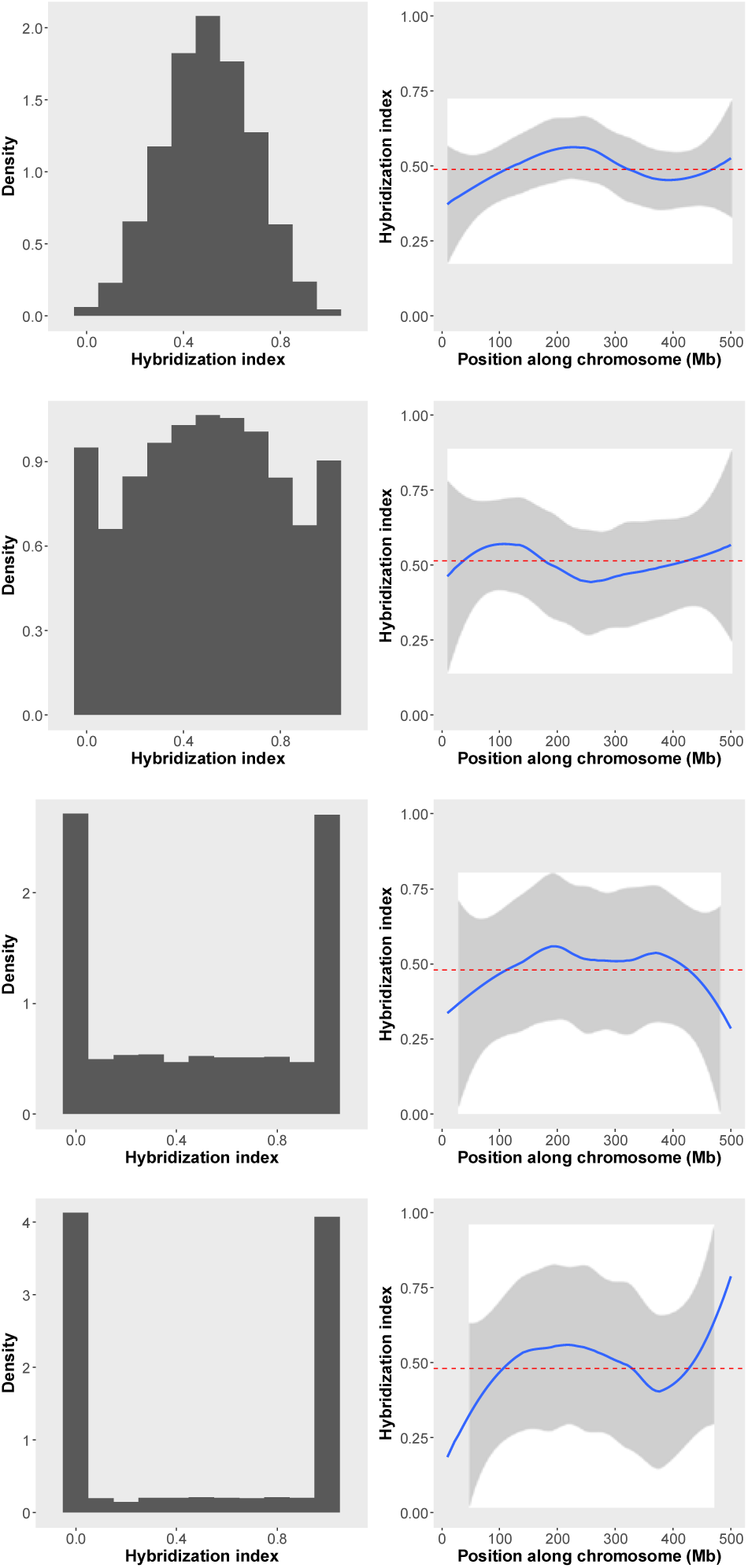
Simulation of the distribution of the hybridization index with various mean size of genomic blocs: from top to bottom: 2, 5, 10, 20 and 50 Mb.

## Supporting information

**Text S1: Jointly inferring phylogenetic relationships and hybridization events**

**Text S2: Determination of possible hybridization**

**Text S3: Inferring multiple hybridizations in four-taxon scenarios**

**Supplementary File 1: *Mathematica* script**

